# Network-based hub biomarker discovery for glaucoma

**DOI:** 10.1101/2022.10.09.511456

**Authors:** Xueli Zhang, Shuo Ma, Xianwen Shang, Xiayin Zhang, Lingcong Kong, Ha Jason, Yu Huang, Zhuoting Zhu, Shunming Liu, Katerina Kiburg, Danli Shi, Yueye Wang, Yining Bao, Hao Lai, Wei Wang, Yijun Hu, Ke Zhao, Guang Hu, Huiying Liang, Honghua Yu, Lei Zhang, Mingguang He

## Abstract

Glaucoma is an optic neuropathy, and the leading cause of irreversible blindness worldwide. However, the early detection of glaucoma remains challenging as chronic forms of glaucoma remain largely asymptomatic until considerable irreversible visual field deficits have ensued. Thus, biomarkers that facilitate early diagnosis and treatment for patients with a high risk of progression are critical. Network medicine approaches can be useful in identifying key relationships and important biomolecules for complex diseases. In this paper, we identified several hub biomarkers/drug targets for the diagnosis, treatment and prognosis for glaucoma and explored their associations for glaucoma based on human disease-biomarker and disease-target-drug networks. These results were verified by text-mining and genomic/epidemiology data. We also predicted the new application of BMP1 and MMP9 to diagnose glaucoma and confirm the theory of hub biomarkers with multiple clinical applications. Further, relevant pivotal pathways (regulation of the multicellular organismal process, regulation of localisation, and cytoplasmic vesicle for biomarkers; signal transduction and developmental process for targets) for these hub biomolecules were discovered, which may be foundations for future biomarker and drug target prediction for glaucoma. In conclusion, based on complex networks, hub biomolecules, essential pathways, and close diseases were identified for glaucoma in diagnosis, treatment and prognosis.

## 1. Introduction

Glaucoma is one of the leading causes of irreversible blindness globally, characterised by retinal ganglion cell loss, retinal nerve fibre layer thinning, and optic disc cupping (1). Glaucoma affects 80 million people and is undiagnosed in nine of ten affected people worldwide (1). Despite the considerable health burden attributable to glaucoma, the accurate diagnosis and treatment of glaucoma remain far from optimal (2–4); indeed, elevated intraocular pressure is the only risk factor that can be treated with medications, laser procedures or glaucoma surgery at present (5). Further research to identify biomarkers critical to the early diagnosis and appropriate therapy for glaucoma is imperative in alleviating its associated burden of disease (6).

The pathogenesis of glaucoma is mediated by multiple genes, while the encoded biomolecules have cross-interactions that are best studies as networks. (7). A wide body of literature has attempted to elucidate the clinical correlations between genetic information, biomarkers and the pathogenesis of glaucoma. Several genes (e.g., *CDKN2B-AS1*, *CAV1* and *CAV2*, *TMCO1*, *ABCA1*, *AFAP1*, *GAS7*, *TXNRD2*, *ATXN2*, *SIX1* and *SIX6*) associated with quantitative glaucoma-related traits (e.g., intraocular pressure, central corneal thickness, and optic disc size) have been identified in genome-wide association studies (8–18). However, there is yet to be one clear pathway that best characterises this. It is likely the multifactorial pathogenesis of glaucoma best lends itself to network studies.

Besides intraocular pressure, oxidative stress, systemic and ocular vascular factors, elevated glutamate concentration and nitric oxide levels, or an autoimmune process have also been implicated in glaucoma pathogenesis (1,19–24). Many biomarkers for glaucoma have been identified, including crystallins, heat shock protein 60 (HSP 60) and HSP 90, myotrophin, apolipoprotein B and apolipoprotein E, endothelial leukocyte adhesion molecule-1, myoblast determination protein 1, myogenin, vasodilator-stimulated phosphoprotein, ankyrin-2 and transthyretin (25–28). Additionally, growing body of evidence supports targets including oxidative stress, and the neuroinflammation might hold great potential for the treatment of glaucoma (29,30). However, given the heterogeneity of experiment samples and environments, it is hard to compare the statistical power and clinical effect for different biomarkers from different studies.

With the advent of the big data era, massive amounts of biomedical data are accumulated and stored on databases. For example, the String database includes the interaction relationships of 19257 human proteins (31), the Therapeutic Target Database (TTD) contains the interaction information between 1512 human diseases to 37316/3419 drugs/targets and 1313 biomarkers(32). These interaction information has helped many studies to detect the relationships for specific biomolecules and explain their biological function. Also, integrated with the interaction information stored in String and TTD, new interactions and functions for biomolecules have been predicted and applied. Hence, it is feasible to integrate the human disease-biomarker information and disease-target-drug information as networks to study them systematically. Our study aimed to fill the gap of analysing glaucoma biomarkers and drug targets based on complex biological interaction networks and found several hub (important points on networks) biomarkers and drug targets. We further identified hub pathways (important pathways) based on these hub biomarkers and drug targets, guiding future biomarkers and drug targets discovery for glaucoma. In addition, the multimorbidity network associated with glaucoma was fully assessed to provide possible underlying biomarkers and pathways.

## 2. Materials and methods

Figure 1 outlines the analysis pipeline for this study.

**Figure 1.**
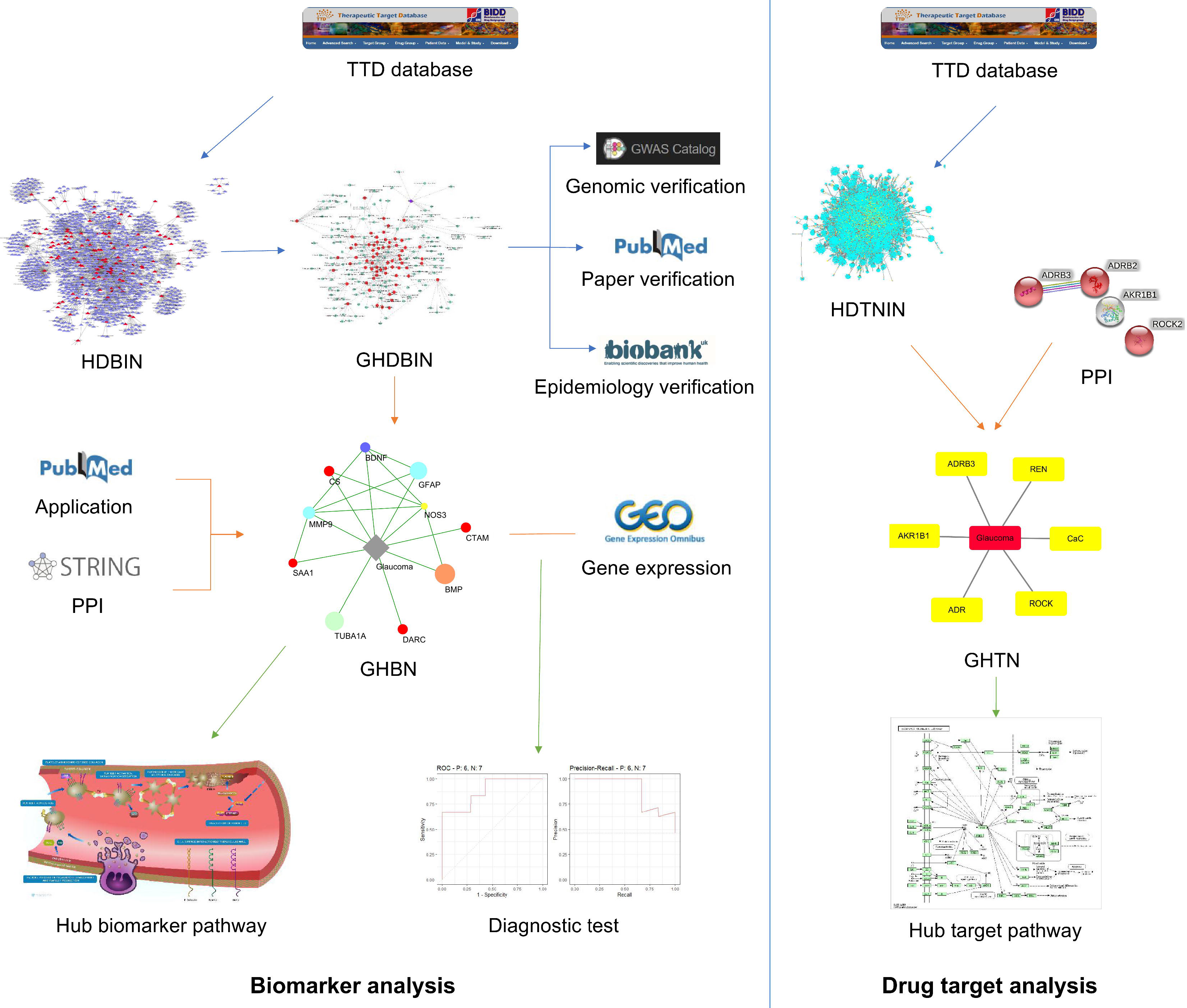
Study pipeline. This study contained 2 parts: the biomarker analysis and the drug-target analysis. The biomarker analysis came from downloading and integrating biomarkers-diseases interaction information to construct the Human disease-biomarker interactions network (HDBIN). Then the greedy search was implemented to find the glaucoma related hub diseases-biomarkers interaction network (GHDBIN). Several close diseases were found on the GHDBIN, and we used GWAS data from GWAS Catalog, text data from PubMed and Epidemiology data from UKB to verify the result. Based on the GHDBIN, we extracted the glaucoma hub biomarker network (GHBN). Further, the application information collected from PubMed and protein-protein interaction (PPI) information from String were added to the hub biomarker network. Gene expression data from GEO was used to verify the diagnosis value of hub biomarkers. Further, biological function analysis was conducted to find key pathways for glaucoma hub biomarkers. The drug-target analysis started with constructing the Human disease-target-drug interactions network (HDTDIN), and the glaucoma hub targets network (GHTN) was selected by the greedy form HDTDIN. PPI information was also annotated to GHTN and critical pathways for glaucoma drug tragets were identified.

### 2.1 Data acquisition

Biomarker-disease interaction information, Drug-target interaction information, target-disease interaction information were downloaded from the TTD database (http://idrblab.net/ttd/).

Protein-protein interaction (PPI) information and pathway information were collected from the String database (https://string-db.org/), the KEGG database (https://www.genome.jp/kegg/), the Reactom database (https://reactome.org/) and the Gene Ontology database (http://geneontology.org/).

Disease related GWAS data were downloaded from the GWAS Catalog database (https://www.ebi.ac.uk/gwas/).

The UK biobank database (https://www.ukbiobank.ac.uk/) provided epidemiological data regarding the patients situation.

Gene expression data were obtained from the GEO database (https://www.ncbi.nlm.nih.gov/geo/, GSE2378, GPL8300 platform).

### 2.2 Construction of Complex Networks

#### Human disease-biomarker interactions network (HDBIN)

We integrated and normalised the human diseases and biomarkers information as standard interaction format in the graph. Based on their interactions, we created a network which we designated as the human disease-biomarker interaction network (HDBIN). Using Cytoscape 3.8.2 software, diseases and biomarkers were marked as points and their relationships as lines.

#### Human disease-target-drug interactions network (HDTDIN)

Meanwhile, the normalised human diseases, drugs, and drug targets and their interaction information were reorganised as a human disease-target-drug interaction network (HDTDIN) by Cytoscape.

### 2.3 Searching for Hub networks

The ‘greedy search algorithm’, a clustering method that takes the best or optimal choice in the current state in each step of the selection, on the *jActiveModules* plugin of Cytoscape was used to find the most active network models (hub networks: the most active small network clustered from the big network) for glaucoma based on HDBIN and HDTDIN. Seven network topology features were selected in searching, and they were: Topological coefficient, Average shortest path length, Closeness centrality, Clustering coefficient, Radiality, Betweenness centrality and Neighbourhood connectivity. In greedy search cluster, the number of sub-models was set as 1000, and the overlap threshold was 0.8. The *jActiveModules* plugin calculated the active levels of clustered networks, and we selected the glaucoma belonged active model as the glaucoma-related hub diseases-biomarkers interaction network (GHDBIN).

The *Pesca* plugin on Cytoscape was then used to calculate the shortest paths from other diseases to glaucoma on the GHDBIN, to find the diseases close to glaucoma on the network. We have hereafter termed these diseases as glaucoma network neighbour diseases (GNND).

### 2.4 Glaucoma related diseases relationships verification test on GWAS data

In order to identify the gene-level relationships for glaucoma and its closest diseases on HNDBIN (network distance very close to glaucoma) found on GHDBIN, GWAS data was used to find the genes that were co-related to glaucoma and other diseases (designated hereafter as inter-genes). Biological network and function analysis was subsequently conducted for these co-specific genes to explore their biological relationships and identify key pathways.

### 2.5 GNND verification test on epidemiological data

The GNND on HNDBIN were verified using the the UK Biobank on epidemiological level. The UK Biobank (UKB) is a population-based cohort of more than 500,000 participants aged 40-73 years who attended one of 22 assessment centers throughout the United Kingdom between 2006 and 2010. Odds ratios (OR) and 95% confidence intervals for diseases associated with glaucoma were evaluated using logistic regression model. Data analyses were conducted using SAS 9.4 software for Windows (SAS Institute Inc.).

### 2.6 Hub biomarker analysis

We selected biomarkers from the GHDBIN to test their relationships on the protein-protein interaction (PPI) network. Biological function analysis was conducted to identify pivotal pathways for glaucoma hub biomarkers.

In order to verify their diagnostic value on biological samples, the Wilcoxon test and receiver operating characteristic curve (ROC) test were conducted using gene expression data by R language.

### 2.7 Glaucoma hub drugs-targets network (GHTN) analysis

The glaucoma hub drugs-targets network (GHTN) generated by application of the greedy search algorithm on HDTDIN was further analysed by biological network and function analysis to find the critical pathways for the prediction of glaucoma drug targets.

### 2.8 Biological function analysis

The String database and ClusterProfiler package on the R language conducted KEGG pathway enrichment analysis and Gene Ontology annotation.

The String database also generated UniProt Annotated Keywords analysis and Reactome Pathway enrichment analysis.

## 3. Results

### 3.1 HDBIN and GHDBIN identified network-based close diseases to glaucoma

Human diseases and biomarkers interaction information was integrated to construct the HDBIN. HDBIN contained 1480 nodes and 2512 lines, including 167 Human diseases and 1313 of their related biomarkers. (Figure 2A, Supplementary Table S1) The GHDBIN identified by the greedy search was illustrated in Figure 2B, which contained 68 glaucoma-related diseases and 123 of their related biomarkers. We calculated the shortest paths between other diseases to glaucoma on GHDBIN. (Supplementary Table S2) The 11 GNND (shortest path=2) was shown on Figure 2C. We searched on PubMed to find the research related to glaucoma and these diseases and found that 9 diseases have been reported together with glaucoma. (Figure 2D)

**Figure 2.**
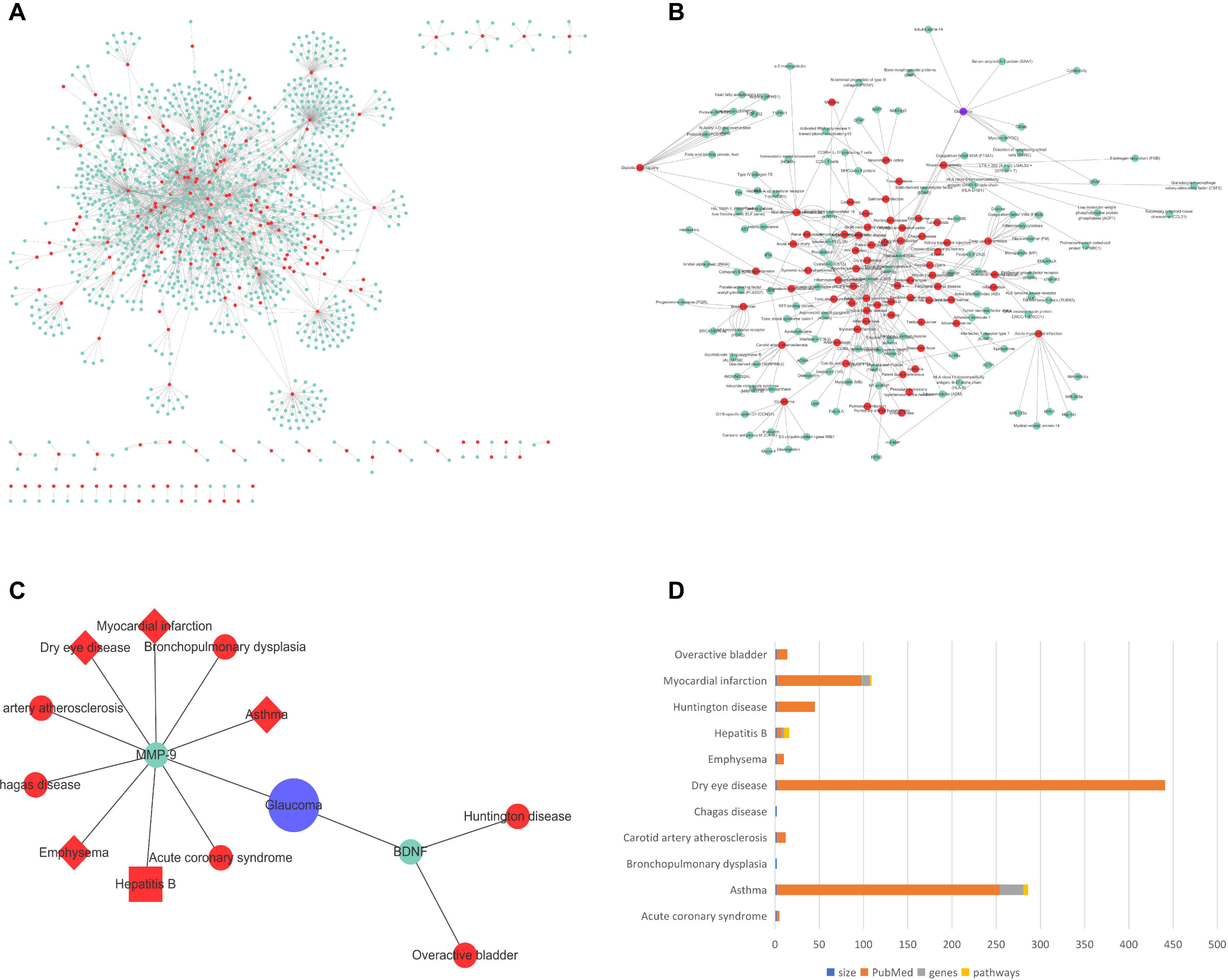
A. Human disease-biomarker interactions network (HDBIN). Red points represented diseases, and green points represented biomarkers. B. Glaucoma related hub diseases-biomarkers interaction network (GHDBIN). The purple point was glaucoma. BDNF and MMP9 were key glaucoma biomarkers connected with other diseases. C. Glaucoma core network on GHDBIN. The shape of points represented a significant level of odd ratio (OR) from UKB data: diamond meant significant and rectangle meant not significant. Point size reflected patients number with glaucoma: the bigger the more. D. Closest diseases to glaucoma on GHDBIN. This histogram reflected the sample size, related paper numbers, number of overlapping genes and pathways for glaucoma and these diseases.

### 3.2 Verification test for GNND on GHDBIN

To facilitate further validation of our findings at a gene level, we downloaded GWAS data for each of the 11 GNND and mapped out the overlap of their corresponding inter-genes. GWAS data for glaucoma and eight other diseases were found, and three diseases (asthma, hepatitis B (HB), and myocardial infarction (MI)) had overlapped specific genes with glaucoma. Supplementary Figure S1 included the LocusZoom plot, which showed the Single Nucleotide Polymorphism (SNP) distribution of GWAS data.

For asthma and glaucoma, 27 inter-genes were identified (Figure 3A), among which only PTHLH-ETS1 and ATXN2-RBFOX1 showed significant interaction in the PPI network. (Figure 3B) Biological function analysis demonstrated that these genes were mapped on five Cellular component pathways: cytoplasmic ribonucleoprotein granule, ribonucleoprotein granule, trans-Golgi network, cytoplasmic stress granule, and histone deacetylase complex. (Figure 3C)

**Figure 3.**
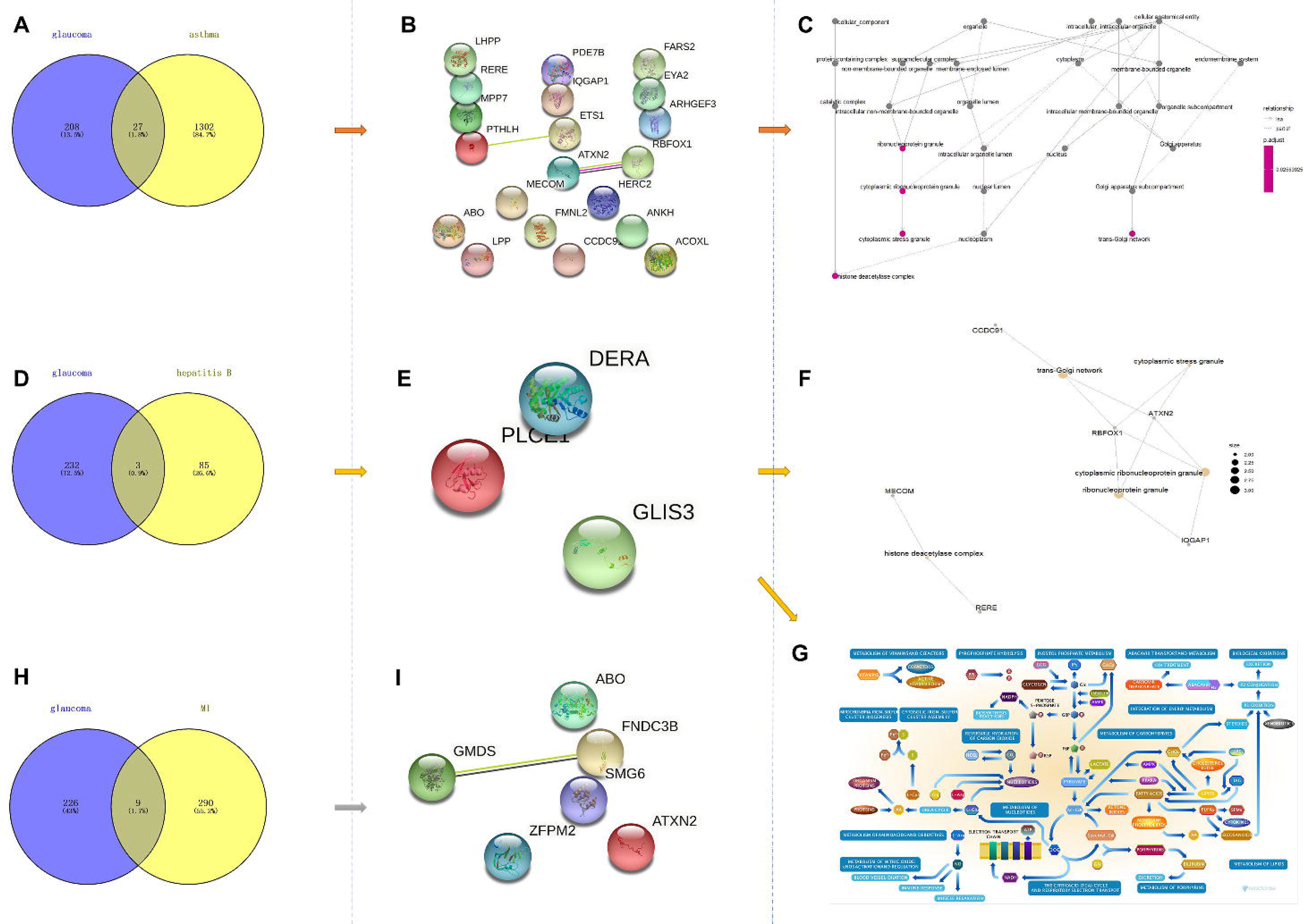
Genomic verification test results for glaucoma closest diseases on GHDBIN. A. There were 27 overlapping genes for glaucoma and asthma from GWAS data. B. PPI network for glaucoma-asthma genes. Only PTHLH-ETS1 and ATXN2-RBFO1 showed significant connections. C. Pathway enrichment analysis results for glaucoma-asthma genes. These genes were mapped mainly on cytoplasmic ribonucleoprotein granule, ribonucleoprotein granule, trans-Golgi network, cytoplasmic stress granule and histone deacetylase complex. D. Glaucoma shared 3 disease-specific genes with hepatitis B. E. No connection was identified for the 3 glaucoma-hepatitis B genes. F. Pathway enrichment results for glaucoma-hepatitis B genes. These genes were mapped on the Metabolism pathway. G. 9 genes were identified as glaucoma-MI shared disease-specific genes. H. PPI network for glaucoma-MI genes. I. GMDS and FNDC3B showed a significant relationship on PPI network.

There were three overlapping genes (DERA, PLCL1, GLIS3) for HB and glaucoma (Figure 3D, 3E), which were not demonstrated to connect on the PPI level. (Figure 3E) GO analysis showed that these genes were associated with trans-Golgi network, cytoplasmic stress granule, cytoplasmic ribonucleoprotein granule, ribonucleoprotein granule, and histone deacetylase complex. Reactome Pathway enrichment analysis showed that these three genes were all related to the Metabolism pathway. (Figure 3G)

Nine genes (CDKN2B-AS1, AC003084.1, AC007568.1, GMDS, ATXN2, ZFPM2, FNDC3B, ABO, and SMG6) overlapped by MI and glaucoma specific genes in GWAS data (Figure 3H), and only GMDS and FNDC3B showed significant connections on the PPI network (Figure 3I). No significant enriched pathways were found for these genes.

Table 1 presented the verification results by the UKB data. There were five closest diseases for glaucoma from GHDBIN found on the UKB, and asthma (OR: 1.10, P-value: 0.028), dry eye disease (OR: 2.16, P-value: 0.00171), and MI (OR: 1.65, P-value: 0) showed significant relationships with glaucoma.

**Table 1.** Verification results on UKB for glaucoma closest diseases on GHDBIN.

|  | Glaucoma |  | Odds ratio (95% CI) | P-value |
| --- | --- | --- | --- | --- |
|  | No | Yes |  |  |
| Asthma |  |  |  |  |
| No | 444231 | 4646 | Reference |  |
| Yes | 58274 | 667 | 1.10(1.01-1.19) | 0.02848 |
| Hepatitis B |  |  |  |  |
| No | 502396 | 5312 | Reference |  |
| Yes | 109 | 1 | 0.87(0.12-6.21) | 0.89361 |
| Emphysema |  |  |  |  |
| No | 502314 | 5309 | Reference |  |
| Yes | 191 | 4 | 2.00(0.74-5.40) | 0.16899 |
| Dry eye disease |  |  |  |  |
| No | 501749 | 5296 | Reference |  |
| Yes | 756 | 17 | 2.16(1.33-3.49) | 0.00171 |
| Myocardial infarction |  |  |  |  |
| No | 491000 | 5117 | Reference |  |
| Yes | 11505 | 196 | 1.65(1.43-1.90) | 0 |

### 3.3 Glaucoma hub biomarkers network (GHBN) presented essential biomarkers for glaucoma

From GHDBIN, we extracted the hub biomarkers (BNDF, CS, MMP9, SAA1, TUBA1A, DARC, BMP, CTAM, NOS3, and GFAP) for glaucoma and integrated them with PPI information, application information (diagnosis, treatment, prognosis) from published papers, and AUCs on diagnostic tests to construct the Glaucoma hub biomarker network (GHBN). (Figure 4A)

**Figure 4.**
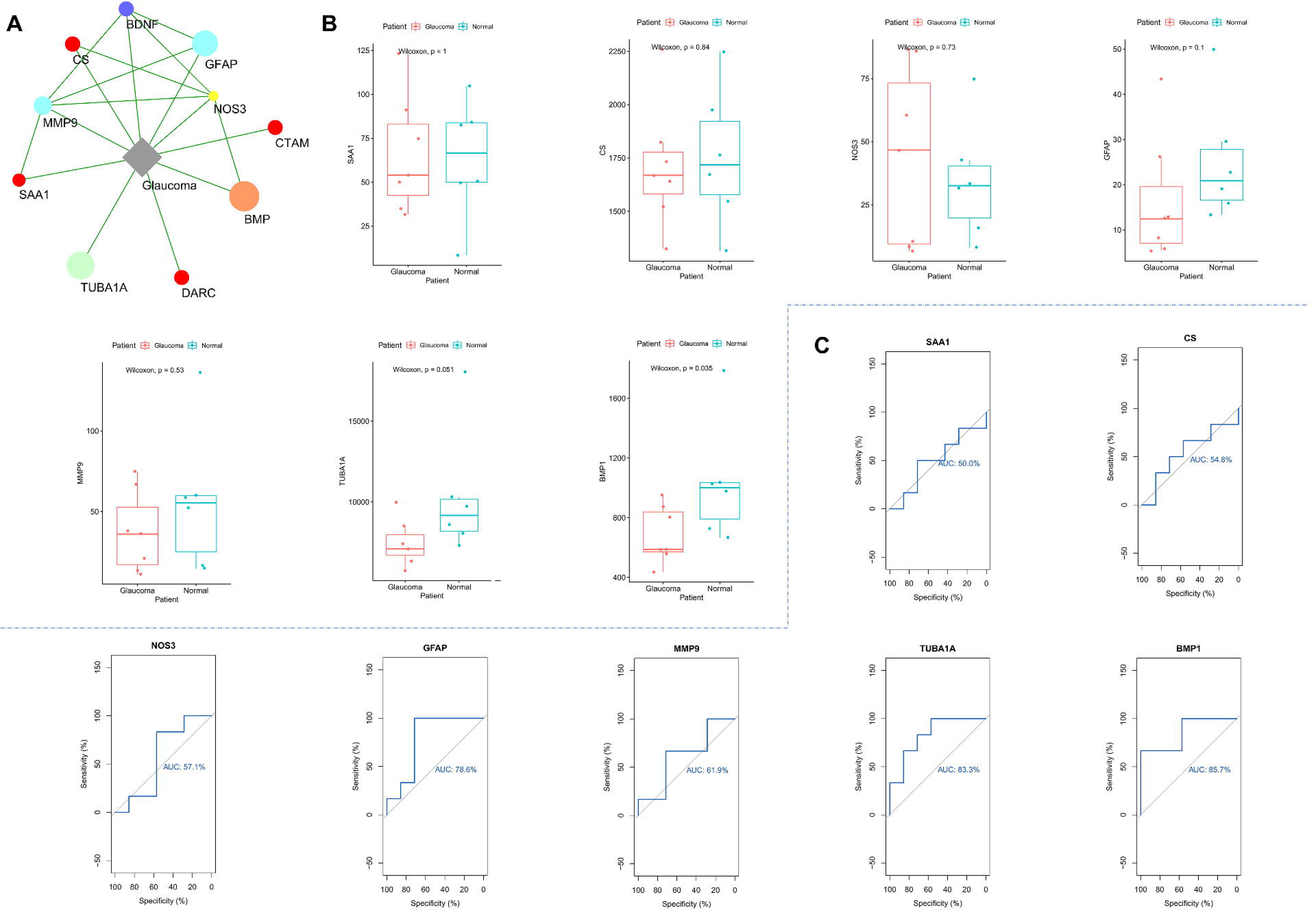
A. Glaucoma hub biomarkers network (GHBN). Different colours represented different applications for these hub biomarkers reported in published papers. Diagnosis/treatment: blue; diagnosis/treatment/prognosis: purple; treatment: red; treatment/prognosis: orange; prognosis: yellow; treatment. B. Expression of hub biomarkers in glaucoma and controls. C. Diagnosis ROC curves for hub biomarkers.

Meanwhile, in order to find common pathways for hub biomarkers, pathway enrichment analysis was conducted. Table 2 presented the significant mapped pathways. On the biological process level, seven biomarkers (BMPER,NOS3,ACKR1,MMP9,SAA1,BDNF, and GFAP) were mapped on the regulation of the multicellular organismal process, and six biomarkers (BMPER,NOS3,MMP9,SAA1,BDNF, and GFAP) were enriched on positive regulation of the multicellular organismal process and regulation of localisation. At the cellular component level, six biomarkers were mapped on the cytoplasmic vesicle, and four were annotated on Methylation by UniProt.

**Table 2.** Biological function analysis results for glaucoma hub biomarkers

| Category | ID | Name | Gene number | FDR |
| --- | --- | --- | --- | --- |
| GO-Biological Process | GO:0051239 | regulation of multicellular organismal process | 7 | 0.01 |
| GO-Biological<br>Process | GO:0051240 | positive regulation<br>of multicellular<br>organismal<br>process | 6 | 0.01 |
| GO-Biological<br>Process | GO:0032879 | regulation of<br>localization | 6 | 0.0278 |
| GO-Biological<br>Process | GO:0045937 | positive regulation<br>of phosphate<br>metabolic process | 5 | 0.01 |
| GO-Biological<br>Process | GO:0010647 | positive regulation<br>of cell<br>communication | 5 | 0.0346 |
| GO-Biological<br>Process | GO:0023056 | positive regulation<br>of signaling | 5 | 0.0346 |
| GO-Biological<br>Process | GO:0001934 | positive regulation<br>of protein<br>phosphorylation | 4 | 0.0349 |
| GO-Biological<br>Process | GO:0010632 | regulation of<br>epithelial cell<br>migration | 3 | 0.0116 |
| GO-Biological<br>Process | GO:1900122 | positive regulation<br>of receptor binding | 2 | 0.01 |
| GO-Cellular<br>Component | GO:0005881 | cytoplasmic<br>microtubule | 2 | 0.013 |
| GO-Cellular<br>Component | GO:0031410 | cytoplasmic<br>vesicle | 6 | 0.013 |
| GO-Cellular<br>Component | GO:0043209 | myelin sheath | 2 | 0.0437 |
| GO-Cellular<br>Component | GO:0055037 | recycling<br>endosome | 2 | 0.0437 |
| GO-Cellular<br>Component | GO:0099513 | polymeric<br>cytoskeletal fiber | 3 | 0.0437 |
| UniProt<br>annotation | KW-0488 | Methylation | 4 | 0.0397 |

### 3.4 Diagnostic test results for glaucoma hub biomarkers

The GSE2378 (platform: GPL8300) gene expression dataset from the GEO database was selected to conduct the diagnostic test for hub glaucoma biomarkers, which included 6 normal and 7 glaucomatous optic nerve head. Figure 4B presented the gene expression comparison between glaucoma and controls in different biomarkers. BMP1 (p value=0.035 in wilcoxon text) showed significant differences between patients and controls.

Diagnostic ROC tests were conducted to verify the diagnostic or prognostic values for these hub biomarkers, of which GFAP (AUC=0.786), MMP9 (AUC=0.619), TUBA1A (AUC=0.833), and BMP1 (AUC=0.857) showed significant diagnostic effects. (Figure 4C) Earlier literature has previously reported the utility of GFAP (5) and MMP9 (33) as diagnostic biomarkers for glaucoma. BMP1 has been reported as both therapeutic (34) and prognostic (35) biomarkers, while TUBA1A has not been previously reported as any glaucoma biomarker. Previous studies have proven that specific biomarkers are likely to perform multiple functions in diagnosing, treating, and prognosis of complex diseases (36). Therefore, we posit that BMP1 and TUBA1A may have future value as diagnostic biomarkers for glaucoma.

### 3.5 HDTDIN identified essential drug targets for glaucoma

HDTDIN contains nodes and 33385 lines, including 1481 Human diseases, 2833 drug targets, and 29071 drugs. (Figure 5A, Supplementary Table S3)

**Figure 5.**
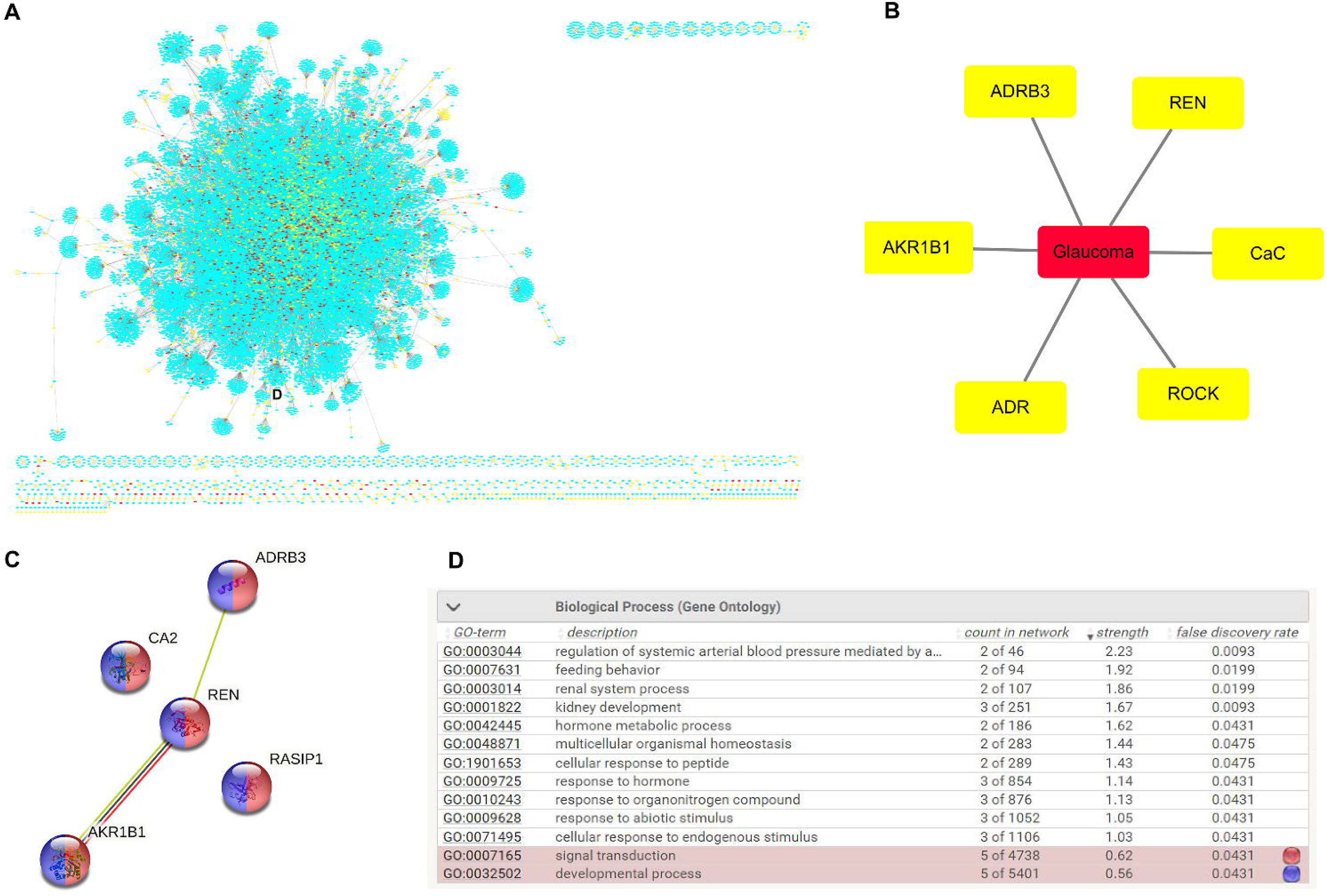
A Human disease-target-drug interactions network (HDTDIN). B. Glaucoma hub drug targets network (GHTN). C. PPI network for glaucoma hub drug targets. D. Biological function analysis results. All 5 protein hub targets were mapped on signal transduction and development process.

The greedy search identified six hub glaucoma drug targets: ADRB3, REN, AKR1B1, CaC, ADR and ROCK, and we constructed the glaucoma hub drug target network. (GHTN) (Figure 5B) We put these targets on the PPI network and found that ADRB3, REN and ADRB2 have been proven with strong relationships. (Figure 5C) Also, biological function analysis found that signal transduction and developmental process on biological process were enriched by all hub protein targets. (Figure 5D)

## 4. Discussion

Along with the development of both traditional and computational experiments, many biomarkers have been reported for glaucoma(37). Our study identified hub glaucoma biomarkers based on HDBIN, which collected all the human biomarkers-diseases interaction information. Then relevant biological function analysis was conducted, and we found several pivotal pathways for these hub biomarkers. These pathways may be foundations for future hub biomarker discovery for glaucoma. Meanwhile, diseases relationships for glaucoma were explored on HDBIN. Further, some hub drug targets and their essential pathways for glaucoma have been identified on HDTDIN. In summary, our study provides key pathways for hub biomarkers/drug targets which are important for clinical-based biomarker/drug discovery in glaucoma.

Analysing diseases and biomarkers in system networks could give a broader view for complex disease research: different from traditional methods, network analysis could investigate disseises situation as a connected system (7). To the best of our knowledge, this is the first study to identify hub biomarkers and related pathways in glaucoma using a network medicine analytical approach. Our study finds 10 glaucoma hub biomarkers, compared with PubMed documentations, five have been used in a single application: GTAM (38), DARC (39) and SAA1 (40) have been used as treatment biomarkers, and CS (41) and NOS3 (42) have been used as prognosis biomarker. Except TUBA1A not found in PubMed, other four hub biomarkers have been used in multiple ways: GFAP (5,43) and MMP9 (33,44) have been applied as diagnosis & treatment biomarkers, BMP (34,35) has been reported as treatment & prognosis biomarkers, and BDNF (45,46) has been reported as diagnosis & treatment & prognosis biomarker. In addition to the biomarkers that have previously been identified in earlier research, we predict that BMP1 and TUBA1A could add their diagnosis ability in the future, which also conforms to our theory that hub biomarkers always conduct multiple uses in complex diseases (47).

Several critical pathways for glaucoma hub biomarkers were identified. No previous studies have reported the definite relationships between glaucoma biomarker or positive regulation of the multicellular organismal process. Hence, we suggest that multicellular organismal process may play a key role in glaucoma and could be a new direction for glaucoma biomarker discovery. Many studies have proven the relationships between glaucoma biomarker with regulation of localisation (48) and cytoplasmic vesicle (49). Four hub biomarkers were enriched on methylation, which add the evidence for DNA methylation is an universal biomarker for complex diseases (50).

We also investigated the disease-disease association relationships on HDBIN in this study. We discovered 11 glaucoma-closest diseases on GHDBIN and verified in genomic and epidemiology data results based on network topology analysis for the first time. Network analysis adds a new feature for detecting disease associations and could widely exclude the heterogeneity of experiments. Further, network relationships could reflect inside disease associations which may not be found by traditional statistical analysis.

Many drugs have been used in the treatment of glaucoma. Based on drug-targets networks, finding the hub drug targets to predict new drugs may be a new strategy for glaucoma drug discovery. This study is the first to identify hub drug targets for glaucoma and found six hub targets: ADRB3, REN, AKR1B1, ADR, ROCK, CaC. These hub targets play important role in biological process: ADRB3 belongs to the beta-adrenergic receptors family, involved in the regulation of fat and thermogenesis. REN participates in regulating blood pressure and electrolyte balance. AKR1B1 is involved in the development of diabetes complications by catalyzing the reduction of glucose to sorbitol. ADR acts as a calcium release channel in the sarcoplasmic reticulum. ROCK plays an important role in cytokinesis. Pathway enrichment analysis found that signal transduction and developmental process were mapped by all five protein hub targets, which may be critical pathways for future glaucoma drug targets prediction. Signal transduction pathway is used to transmit the information of the ligand to the change in the biological activity of the target cell (51). Our study adds evidence for signal transduction as an important platform for drug discovery (52).

The strength of our study is that we firstly identified hub biomarkers/drug targets for glaucoma based on network theory and verified related findings in different datasets. However, our methods and results need to be verified further in other experiments.

## 5. Conclusions

In conclusion, we have used a network medicine analysis to identify novel and validate existing diagnostic and therapeutic biomarkers for glaucoma, and detect relationships between glaucoma and associated diseases. This approach may have value in developing new diagnostic and therapeutic approaches for glaucoma.

## Availability of Data and Material

Network information can be found from the TTD database (http://idrblab.net/ttd/) and the String database (https://string-db.org/).

Pathway information can be fournd on the KEGG database (https://www.genome.jp/kegg/), the Reactom database (https://reactome.org/) and the Gene Ontology database (http://geneontology.org/).

Genomic data can be found on the GWAS Catalog database (https://www.ebi.ac.uk/gwas/) and the GEO database (https://www.ncbi.nlm.nih.gov/geo/).

The UK biobank database (https://www.ukbiobank.ac.uk/) provided epidemiological data concerning the patients situation.

## Author Contributions

**Xueli Zhang**: Conceptualization, Investigation, Data curation, Methodology,Visualization. **Shuo Ma**: Data curation, Formal analysis, Validation, Writing–original draft. **Xianwen Shang**: Validation, Writing–review & editing. **Xiayin Zhang**: Methodology, Writing–review & editing. **Lingcong Kong**: Methodology, Writing–review & editing. **Ha Jason**: Writing– review & editing. **Yu Huang**: Writing–review & editing. **Zhuoting Zhu**: Writing–review & editing. **Shunming Liu**: Writing–review & editing. **Katerina Kiburg**: Writing–review & editing. **Danli Shi**: Writing–review & editing. **Yueye Wang**: Writing–review & editing. **Yining Bao**: Writing–review & editing. **Hao Lai**: Writing–review & editing. **Wei Wang**: Writing–review & editing. **Yijun Hu**: Writing–review & editing. **Ke Zhao**: Writing–review & editing. **Guang Hu**: Writing–review & editing. **Huiying Liang**: Supervision, Writing– review & editing. **Honghua Yu**: Supervision, Writing–review & editing. **Lei Zhang**: Supervision, Writing–review & editing. **Mingguang He**: Conceptualization, Supervision, Writing–review & editing.

## Acknowledgements

The authors are grateful to our colleagues and collaborators, especially Yuanyuan Chen. This research was funded by the Fundamental Research Funds of the State Key Laboratory of Ophthalmology, Zhongshan Ophthalmic Center and the GDPH Supporting Fund for Talent Program (KJ2020633, KJ012019530). Prof. Mingguang He receives support from the University of Melbourne Research Accelerator Program and the CERA Foundation. The Centre for Eye Research Australia (CERA) receives Operational Infrastructure Support from the Victorian State Government. The sponsor or funding organisation had no role in the design or conduct of this research.

## Appendix A. Supplementary data

The following are the Supplementary data to this article:

Download : Download PDF file (123kB)

Supplementary data 1.

Download : Download zip file (1.48MB)

Supplementary data 2.

