## Supplementary material for "Network-based hub biomarker discovery for glaucoma": Figure S1

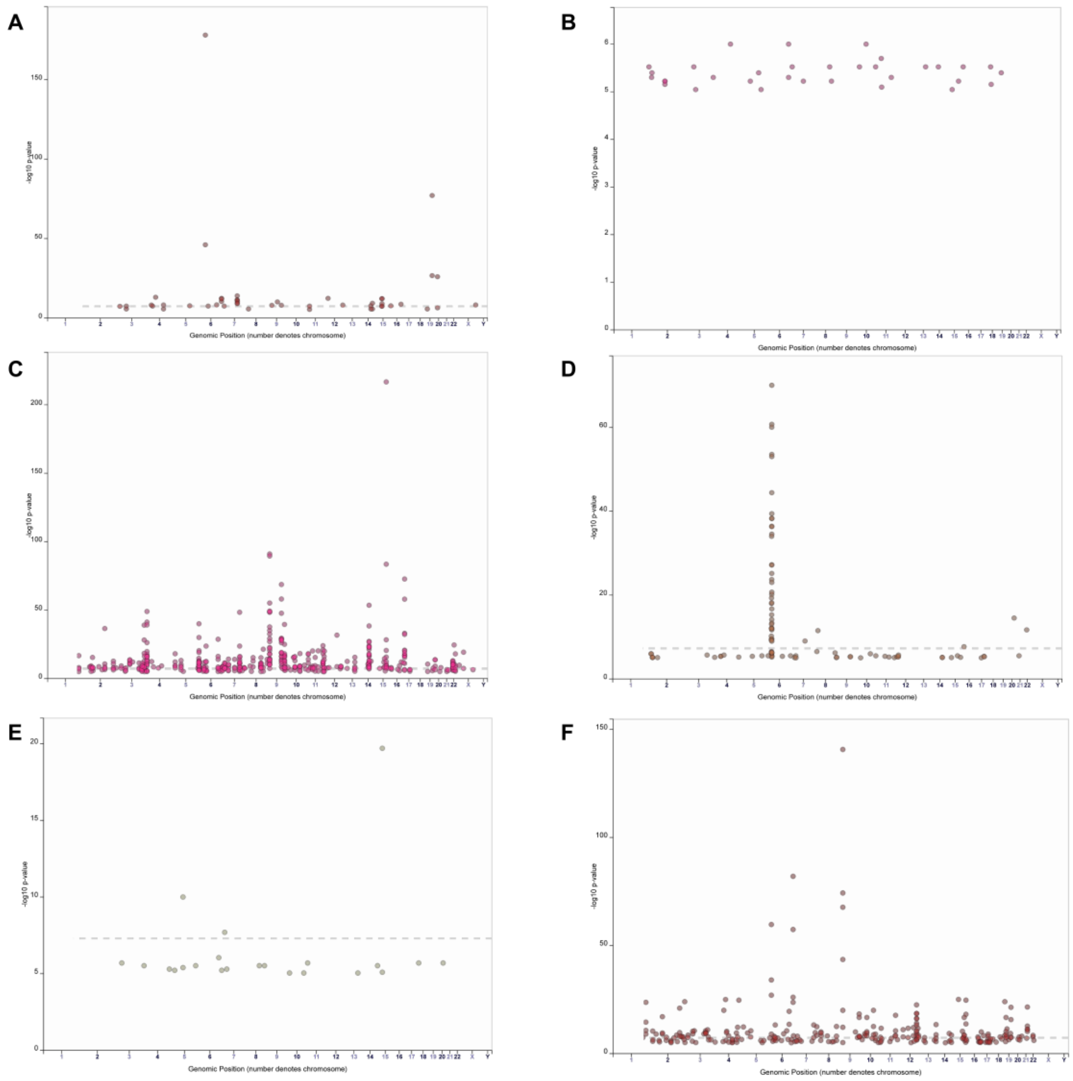


Figure S1. Manhattan plot for GWAS data concerning glaucoma and its related diseases. A. Acute coronary syndrome. B. Bronchopulmonary dysplasia. C. Glaucoma. D. Hepatitis B. E. Huntington disease. F. Myocardial infarction.
